# An introduction to LifeLines DEEP: study design and baseline characteristics

**DOI:** 10.1101/009217

**Authors:** Ettje F. Tigchelaar, Alexandra Zhernakova, Jackie A.M. Dekens, Gerben Hermes, Agnieszka Baranska, Zlatan Mujagic, Morris A. Swertz, Angélica M. Muñoz, Patrick Deelen, Maria C. Cénit, Lude Franke, Salome Scholtens, Ronald P. Stolk, Cisca Wijmenga, Edith J.M. Feskens

## Abstract

There is a critical need for population-based prospective cohort studies because they follow individuals before the onset of disease, allowing for studies that can identify biomarkers and disease-modifying effects and thereby contributing to systems epidemiology. This paper describes the design and baseline characteristics of an intensively examined subpopulation of the LifeLines cohort in the Netherlands. For this unique sub-cohort, LifeLines DEEP, additional blood (n=1387), exhaled air (n=1425), fecal samples (n=1248) and gastrointestinal health questionnaires (n=1176) were collected for analysis of the genome, epigenome, transcriptome, microbiome, metabolome and other biological levels. Here, we provide an overview of the different data layers in LifeLines DEEP and present baseline characteristics of the study population including food intake and quality of life. We also describe how the LifeLines DEEP cohort allows for the detailed investigation of genetic, genomic and metabolic variation on a wealth of phenotypic outcomes. Finally, we examine the determinants of gastrointestinal health, an area of particular interest to us that can be addressed by LifeLines DEEP.

## INTRODUCTION

Many diseases are multifactorial in origin, meaning that they are caused by a combination of genetic and environmental components. To date, a considerable number of genetic variants have been identified that are associated with almost every multifactorial disease or trait.[1] These independent genetic factors are often common, occurring frequently in the absence of disease, and therefore cannot yet be used to predict disease. For example, 40 risk loci have been identified for celiac disease that explain about 54% of disease risk [2], yet there is no clear correlation between carrying these risk alleles and actually developing celiac disease [3]. Thus the question remains: why do some people develop the disease while others are resilient despite carrying many genetic risk alleles? These resilient individuals may provide important clues to disease prevention, but they can only be identified when apparently healthy individuals are followed over time. This highlights the need for prospective cohort studies where life course processes are investigated and determinants of health and disease are identified. An advantage of population-based prospective cohort studies is that they are not specifically targeted to a diseased population and they follow individuals before disease onset, allowing for studies that can identify biomarkers and disease-modifying effects.[4] Furthermore, age-related processes that correlate to health and disease can be studied in these cohorts.

LifeLines is a population cohort of over 165,000 participants which covers multiple generations and focuses on determinants for multifactorial diseases. This large-scale population cohort includes detailed information on phenotypic and environmental factors, change of risk factors, quality of life, and disease incidence.[5] Genetic information is also available for about 10% of the population. A subset of approximately 1500 LifeLines participants also take part in LifeLines DEEP. These participants are examined more thoroughly, specifically with respect to molecular data, which allow for a more thorough investigation of the association between genetic and phenotypic variation. For these participants, additional biological materials are collected, as well as additional information on environmental factors. Subsequently, genome-wide transcriptomics and methylation data are generated, metabolites and biomarkers are measured, and the gut microbiome is assessed.

LifeLines DEEP specifically allows for in-depth analysis of gastrointestinal-health-related problems like irritable bowel syndrome (IBS). This is an important direction for research since gastrointestinal complaints are highly prevalent in the general population and have a high impact on quality of life.[6–8] IBS is a functional bowel disorder which involves abdominal pain or discomfort and related change in bowel habits.[9] Prevalence of IBS in Western countries varies widely among different studies ranging from 4% up to 22%.[10] There are, however, no specific tests available to diagnose IBS. The current diagnosis is based on excluding gastrointestinal diseases and on symptoms using diagnostic criteria such as the Rome III criteria.[9]

In this paper we describe the study design and baseline characteristics of the LifeLines DEEP cohort and explain how the collected data can be applied to multiple fields of interest.

## METHODS

### LifeLines

Individuals aged 25-50 years old were invited by their general practitioner to participate in the LifeLines study. Upon inclusion, participants were asked about their family. Family members were then also invited to participate in LifeLines in order to obtain inclusion over three generations. At baseline, all participants visited one of the LifeLines Research Sites twice for physical examinations. Prior to these visits, two extensive baseline questionnaires were completed at home. At the first visit, anthropometry, blood pressure, cognitive functioning and pulmonary function as well as other factors were measured (Table 1a). At the second visit, approximately two weeks later, a fasting blood sample was collected. In total 167,729 participants have been included who will be followed for 30 years. Every eighteen months, each participant receives a follow-up questionnaire. Additionally, once every five years, each participant is invited to one of the LifeLines Research Sites for follow-up measurement of their health parameters.

**Table 1a.** Overview of collected data in LifeLines and source it originates from. More detailed information is available through the LifeLines data catalog at www.lifelines.net

| **LifeLines** |  |
| --- | --- |
| **Data** | **Source** |
| 24h urine biochemistry | Assessed by trained technician |
| Blood basic chemistry | Assessed by trained technician |
| Anthropometry | Examined by trained physician assistant |
| Blood pressure | Examined by trained physician assistant |
| Cognitive functioning | Examined by trained physician assistant |
| Electrocardiogram | Examined by trained physician assistant |
| Pulmonary function | Examined by trained physician assistant |
| Demographics | Questionnaire |
| Education | Questionnaire |
| Food intake | Questionnaire |
| Health status | Questionnaire |
| Medication use | Questionnaire |
| Physical activity | Questionnaire |
| Quality of life | Questionnaire |
| Smoking | Questionnaire |

### LifeLines DEEP

From April to August 2013, all participants registered at the LifeLines Research Site in Groningen were invited to participate in LifeLines DEEP in addition to the regular LifeLines program. After participants signed informed consent, three additional tubes of blood were drawn by one of the LifeLines physician assistants during the participant's second visit to site. Exhaled air was also collected during this visit and participants were given instructions for feces collection at home by one of the LifeLines DEEP assistants. The participants who agreed to collect a fecal sample also received the questionnaire on gastrointestinal complaints. Immediately after fecal sample collection, the sample was frozen at −20°C. Fecal samples were collected on dry-ice from the participants’ homes within 1 to 2 weeks after the second site visit. Upon arrival at the research location, fecal samples were immediately stored at −80°C.

### Inclusion

Initially, 1539 participants were included in the LifeLines DEEP study. Of these participants, 78 dropped-out: 51 did not complete the second visit to the LifeLines location in time and 27 withdrew from participation. In total, 1461 individuals completed the LifeLines DEEP study. From those participants, we collected additional blood for genetics, methylation, and transcriptomics analyses (n=1387); exhaled air for analysis of volatile organic compounds (n=1425); and fecal samples for microbiome and biomarker assessment (n=1248) (Table 1b). Moreover, 1176 gastrointestinal complaints questionnaires were returned. For 1183 participants, we collected all three biomaterials: additional blood, exhaled air and feces (Fig 1). For 168 participants, we collected additional blood and exhaled air and for 65 participants we collected exhaled air and feces. For 45 participants, we only have additional blood or exhaled air.

**Table 1b.** Overview of additional data collected in LifeLines DEEP including the number of samples, the source biomaterial it originates from and the method of analysis used

| **LifeLines DEEP** |  |  |  |
| --- | --- | --- | --- |
| **Data** | **n** | **Source** | **Method** |
| Biological aging | 1387 | Cells from whole blood | FlowFish |
| Biological aging | 1387 | DNA from whole blood | qPCR and sjTRECs |
| Biomarkers<br>(citrulline, cytokines) | 1387 | Plasma | HPLC, ECLIA |
| Biomarkers<br>(calprotectin, HBD-2,<br>chromogranin A, SCFA) | 1248 | Feces | ELISA, ELISA,<br>RIA, GC-MS |
| CVD risk score | 1448 | Biochemical measures<br>and questionnaire | Scoring algorithm<br>Framingham Heart Study |
| Functional studies | 1387 | PBMC from whole blood | Various methods |
| Gastrointestinal<br>complaints | 1176 | Questionnaire | Rome III criteria and Bristol<br>Stool Form Scale |
| Genetics | 1387 | DNA from whole blood | CytoSNP and ImmunoChip,<br>GONL as imputation reference |
| Metabolomics | 1425 | Exhaled air | GC-tof-MS |
| Metabolomics | 1387 | Plasma | NMR |
| Methylation | 761+ | DNA from whole blood | 450K chip |
| Microbiome | 1248 | Feces | 16S rRNA based sequencing |
| Transcriptomics | 1387 | Whole blood (PAXgene) | RNA sequencing |

*qPCR = quantitative polymerase chain reaction, sjTRECs = signal joint T cell receptor excision circles, HPLC = high-performance liquid chromatography, ECLIA = electro-chemiluminescence immunoassay, HBD-2 = human $\beta$ defensin 2, SCFA = short chain fatty acids, ELISA = enzyme-linked immunosorbent assay, RIA = radioimmunoassay, GC-(tof-)MS = gas chromatography-(time of flight-)mass spectrometry, NMR = nuclear magnetic resonance*

**Fig 1.**
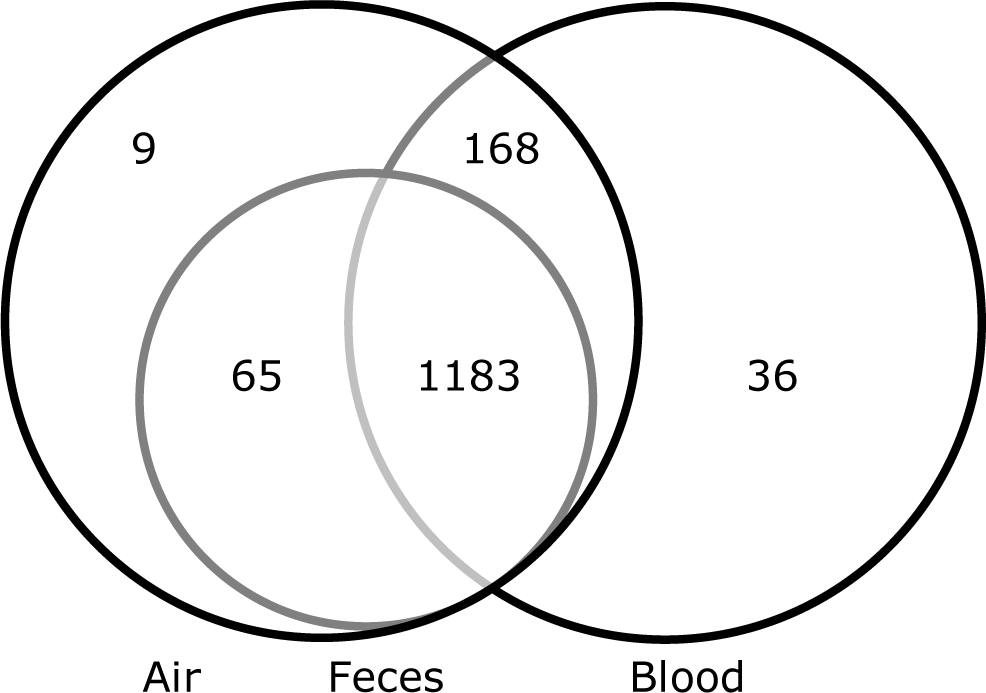
Overview of additional biomaterials collected in LifeLines DEEP

## Additional data types

Genome wide *transcriptomics* were assessed as a measure of gene expression. We isolated RNA from whole blood collected in a PAXgene tube using PAXgene Blood miRNA Kit (Qiagen, CA, USA). The RNA samples were quantified and assessed for integrity before sequencing. Total RNA from whole blood was deprived of globin using GLOBINclear kit (Ambion, Austin, TX, USA) and subsequently processed for sequencing using Truseq version 2 library preparation kit (Illumina Inc., San Diego, CA, USA). Paired-end sequencing of 2 x 50bp was performed using Illumina's Hiseq2000, pooling 10 samples per lane. Finally, read sets per sample were generated using CASAVA, retaining only reads passing Illumina's Chastity Filter for further processing. On average, the number of raw reads per individual after QC was 44.3 million. After adapter trimming, the reads were mapped to human genome build 37 using STAR (https://code.google.com/p/rna-star/). Of these, 96% of reads were successfully mapped to the genome. Transcription was quantified on the gene and meta-exon level using BEDTools (https://code.google.com/p/bedtools/) and custom scripts, and on the transcript level using FluxCapacitor (http://sammeth.net/confluence/display/FLUX/Home).

We isolated total DNA from EDTA tubes and profiled genome wide *methylation* using the Infinium HumanMethylation450 BeadChip, as previously described.[11] In short, 500 ng of genomic DNA was bisulfite modified and used for hybridization on Infinium HumanMethylation450 BeadChips, according to the Illumina Infinium HD Methylation protocol.

We determined *metabolites* in both exhaled air and blood. Metabolites from exhaled air were measured by a combination of gas chromatography and time-of-flight mass spectrometry (GC-tof-MS) as described previously[12,13]. In short, the exhaled air sample was introduced in a GC that separates the different compounds in the mixture. Subsequently, the compounds were introduced into the MS to detect and also to identify the separated volatile organic compounds. The metabolites in plasma were measured using the nuclear magnetic resonance (NMR) method, as described by Kettunen et al.[14]

*Genotyping* of genomic DNA was performed using both the HumanCytoSNP-12 BeadChip [15] and the ImmunoChip, a customized Illumina Infinium array [16]. Genotyping was successful for 1,385 samples (CytoSNP) and 1,374 samples (IChip), respectively. First, SNP quality control was applied independently for both platforms. SNPs were filtered on MAF above 0.001, a HWE p-value >1e^−4^ and call rate of 0.98 using Plink.[17] The genotypes from both platforms were merged into one dataset. For genotypes present on both platforms the genotypes were put on missing in the case of non-concordant calls. After merging, SNPs were filtered again on MAF 0.05 and call rate of 0.98 resulting in a total of 379,885 genotyped SNPs. Next, this data was imputed based on the Genome of the Netherlands (GoNL) reference panel.[18–20] The merged genotypes were pre-phased using SHAPEIT24[21] and aligned to the GoNL reference panel using Genotype Harmonizer (http://www.molgenis.org/systemsgenetics/) in order to resolve strand issues. The imputation was performed using IMPUTE2[22] version 2.3.0 against the GoNL reference panel. We used MOLGENIS compute [23] imputation pipeline to generate our scripts and monitor the imputation. Imputation yielded 8,606,371 variants with Info score ≥0.8. In addition, HLA type was established via Broad SNP2HLA imputation pipeline.[24]

We collected several types of *cells* including lymphocytes and granulocytes for assessment of telomere length as a measure for aging. Currently, we are optimizing the FlowFish method of telomere measuring as described by Baerlocher et al.[25] In addition, peripheral blood mononuclear cells (PBMCs) were collected and stored at −80°C for future functional studies.

Fecal samples were collected in order to study the *gut microbiome*. Gut microbial composition was assessed by 16S rRNA gene sequencing of the V4 variable region on the Illumina MiSeq platform according to the manufacturer's specification.[26] Reads were quality filtered and taxonomy was inferred using a closed reference Operational Taxonomic Unit picking protocol against a pre-clustered GreenGenes database, as implemented by QIIME.[27,28] Moreover, fecal aliquots were stored for future analysis of gastrointestinal– health-related biomarkers.

In addition, phenotypic data was collected on *gastrointestinal health complaints* by means of the Rome III criteria questionnaire [9] and the Bristol Stool Form Scale.[29]

We collected and stored *plasma* for future analysis of disease and ageing-related biomarkers such as circulating microRNAs.

### Analyses of baseline characteristics, quality of life, gastrointestinal complaints and qualitative food intake

For each participant a risk score for cardiovascular disease (CVD) was calculated according to the scoring algorithm developed within the Framingham Heart Study.[30] The CVD risk score ranges from ≤-3 to ≥18 and is calculated based on gender, age, HDL, total cholesterol, systolic blood pressure, smoking status, and presence or absence of diabetes.

We calculated quality of life scores based on the RAND 36-item Short Form Health Survey scoring version I by calculating eight summary scores and the mental and physical component score.[31–33] The summary scores range from zero to 100 points where 100 represents the best quality of life. The mental and physical component scores were transformed to have a mean of 50 and a standard deviation of 10 compared to the reference population as described by Ware et al.[32–34]

Occurrence of functional bowel disorders was assessed via the Rome III criteria.[9] Participants with self-reported Crohn's disease, ulcerative colitis and celiac disease were excluded from this analysis.

Data on habitual dietary intake was collected via a validated food frequency questionnaire, developed by the division of Human Nutrition of Wageningen University.[35]

Mean and standard deviations for the baseline characteristics and quality of life scores were calculated. Statistical programs R (version 3.0.1) and IBM SPSS Statistics (version 20) were used for analyses and construction of figures.

## RESULTS

Over a period of 6 months, 1539 participants enrolled in the LifeLines DEEP study. Slightly more women (n=903, 58.7%) than men (n=636, 41.3%) were included (Table 2). The age of the participants ranged from 18 to 86 years, with a mean age of 44. Mean BMI was 25.2 kg/m^2^. On average, total cholesterol level and blood glucose level both were 5.0 mmol/L. Average blood pressure was lower in women (116/68 mmHg) compared to men (124/74 mmHg). Amongst women, the percentage of current smokers was a little lower (18.3%) than in men (19.5%). The Framingham risk score for cardiovascular disease was on average 5.7 for women and 8.6 for men, corresponding to a 3% and 7% risk of a first cardiovascular event, respectively (Table 2).[30] In our cohort the quality of life score was lowest for vitality (mean(sd): 67.2(15.5)) and highest for physical functioning (mean(sd): 92.1(12.3)) (Fig. 2 and Supplementary table Table 3). The age-and gender-adjusted quality of life scores yielded similar results.

**Fig 2.**
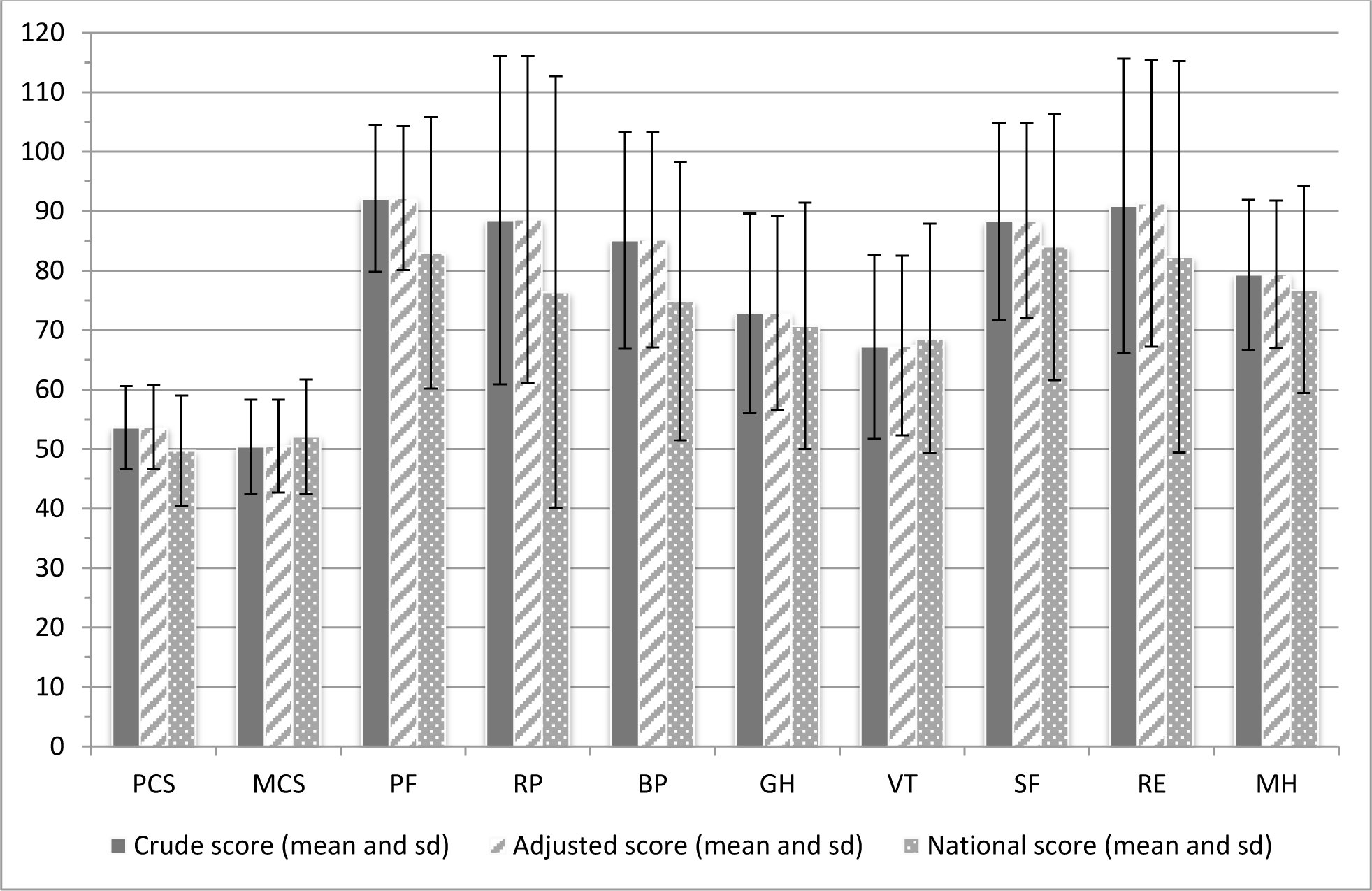
Mean and sd of crude and adjusted quality of life scores, two component scores and eight group scores in the LifeLines DEEP population (n=1539) compared to a national sample of the Dutch population [34,42]. *Adjusted score is adjusted for gender and age. PCS: physical component score, MCS: mental component score, PF: physical functioning, RP: role-physical, BP: bodily pain, GH: general health, VT: vitality, SF: social functioning, RE: role-emotional, MH: mental health*.

**Table 2.**
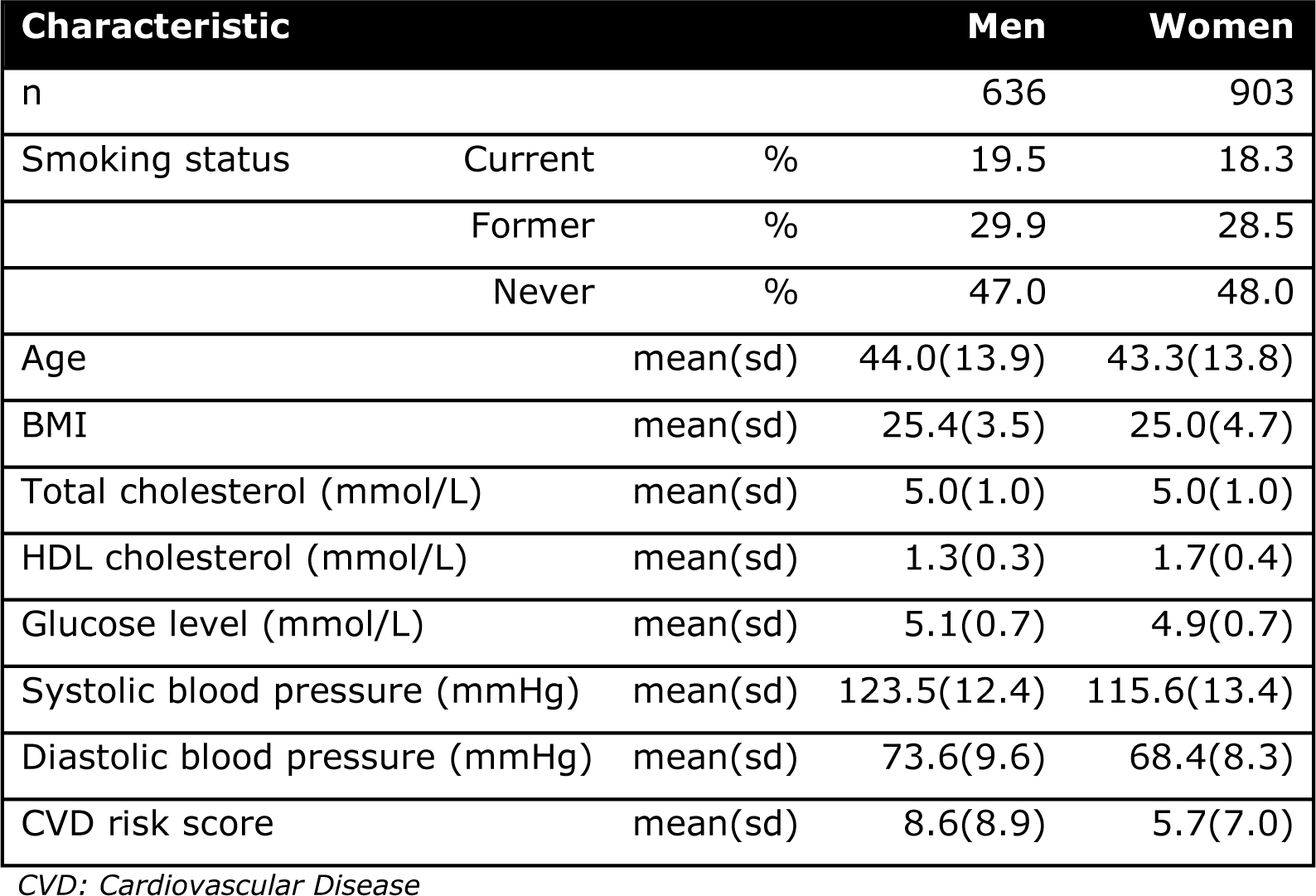
Baseline characteristics LifeLines DEEP by gender including smoking, age, BMI, cholesterol level, glucose level, blood pressure and Framingham risk score for cardiovascular disease

Analysis of 1176 gastrointestinal complaints questionnaires identified 409 participants with functional bowel disorders (Fig 3). Prevalence of irritable bowel syndrome (IBS) in our cohort was 21% (n=249). Another 13% (n=160) of participants fulfilled criteria for functional bloating (9%, n=108) or functional constipation (3%, n=37) or functional diarrhea (1%, n=15). Two third of the participants (n=767) did not meet the Rome III criteria for functional bowel disorders. Moreover, 4% (n=51) of the participants answered that they never experience any gastrointestinal complaints (Fig 3).

**Fig 3.**
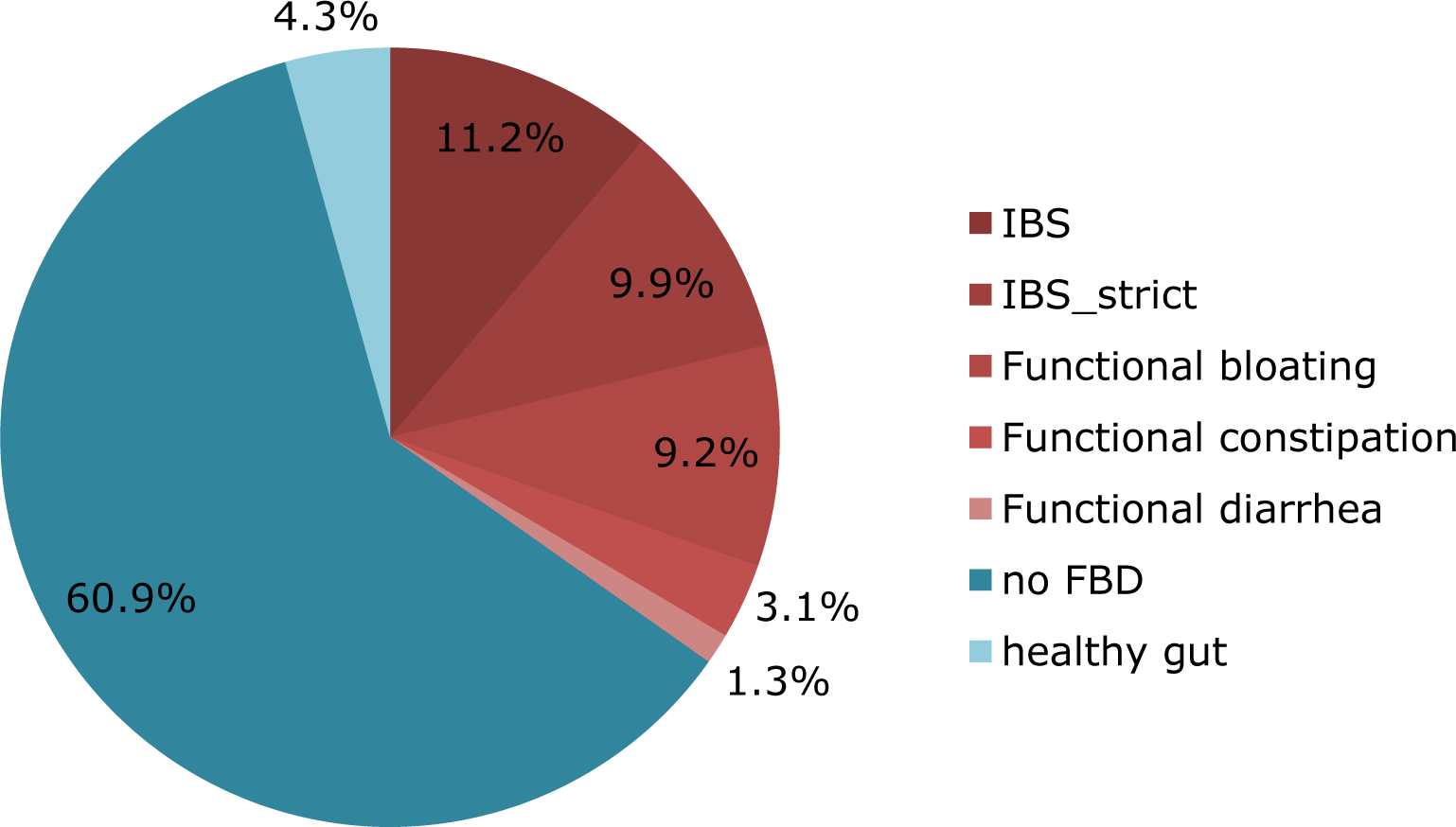
Functional bowel disorders in the LifeLines DEEP cohort based on Rome III criteria (n=1176) *IBS (Irritable Bowel Syndrome): pain or discomfort at least 2-3 days/month, IBS_strict: pain or discomfort more than one day per week, FBD: functional bowel disorder, healthy gut: lowest possible score on Rome III questionnaire*

Analysis of the frequency of intake of major food groups showed a subdivision into three main categories. The first category contained food groups that were consumed daily, bread and coffee, for example (Fig 4a and 4b). The second category contained food groups for which consumption ranged from daily to a few days per week. Examples of these food groups include meat, vegetables and fruit (Fig 4c, 4d and 4e). The third category included food groups that were consumed on a weekly to monthly basis only, fish, for example (Fig 4f). For other food groups, such as milk and alcoholic beverages, the intake varied greatly (Fig 3g and 3h). This frequency data will later be combined with portion sizes and the Dutch food composition table [NEVO 2006, RIVM, Bilthoven] to estimate nutrient intake in grams per day.

**Fig 4.**
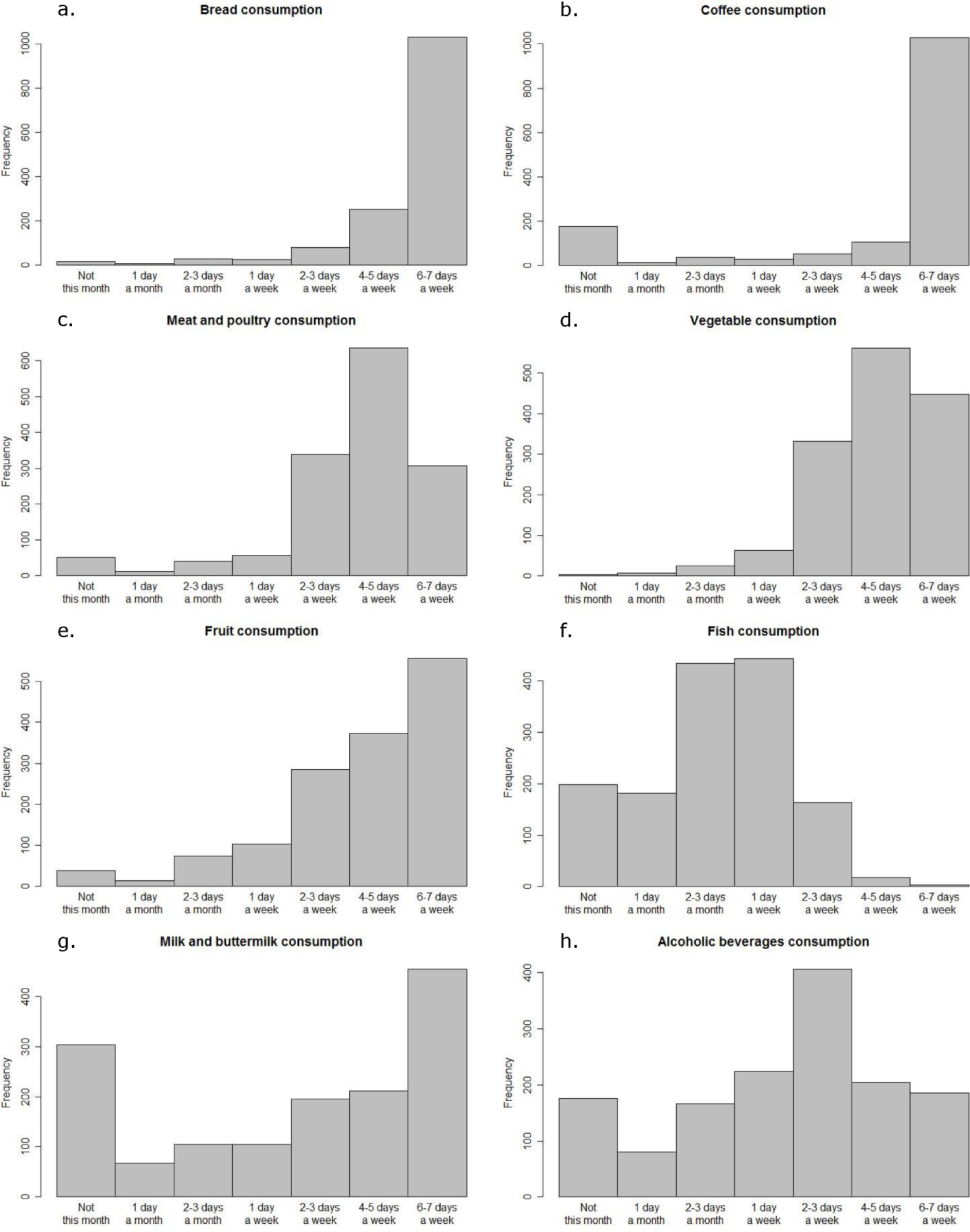
Qualitative intake of a. bread, b. coffee, c. meat and poultry, d. vegetables, e. fruit, f. fish, g. milk and buttermilk, and h. alcoholic beverages in LifeLines DEEP (n=1539). Bars represent: ‘not this month’, ‘1 day a month’, ‘2-3 days a month’, ‘1 day a week’, ‘2-3 days a week’, ‘4-5 days a week’, and ‘6-7 days a week’.

For all individuals, additional biomaterials were collected (Fig 1) for future systems epidemiological studies [36] integrating multi-level ‘omics’ data with environmental, physical and epidemiological data to provide a deeper and more detailed view of the LifeLines DEEP population. These biomaterials include plasma to examine the concentration of metabolites, peripheral blood mononuclear cells to determine genome-wide transcription and methylation profiles, exhaled air to analyze volatile organic compounds and feces to establish composition of the gut microbiome. Moreover, genetic data has been generated for all individuals allowing for the construction of genetic risk profiles for a wide variety of common diseases. These multiple data levels will provide rich opportunities for future research into the molecular underpinnings of human health and disease, as well as research into the interaction between molecular and environmental components including behavior, socio-demographic factors, and analysis of specific subgroups.

## DISCUSSION AND FUTURE PERSPECTIVES

In the design of a population cohort study it is important to balance breadth (the number of samples included) and depth (the amount of phenotypic data). LifeLines is a large prospective cohort which includes more than 165,000 individuals and measures several thousands of phenotypes ranging from biochemical parameters, physical measurements, psychosocial characteristics and environmental factors to detailed information on health status. However, the cohort was not set up to include molecular data levels for the study of health and disease in human populations. With LifeLines DEEP, we are performing a pilot study of additional deep molecular measurements in 1,500 individuals using biomaterials from different domains that were all collected contemporaneously from fasting individuals. Although the LifeLines DEEP cohort is relatively small, it will allow for proof-of-concept studies into systems epidemiology. LifeLines DEEP is unique in having exhaled air measurements from all individuals and a level of information that, to our knowledge, is rarely present in other population-based cohorts. Additionally, both the collection of cells for telomere length measurements and further functional studies and the fecal sample collection are unique. LifeLines DEEP will not only contribute to a better understanding of the association between genetic variation and molecular function, but can also be integrated with other population cohorts that have similar molecular data. In particular, the collection and analysis of fecal material is crucial given increasing evidence that the gut microbiome can play an important role in health and disease [37]. Nevertheless, harmonization and linking of data across multiple cohorts might be needed to achieve critical numbers. The Biobanking and Biomolecular Research Infrastructure in the Netherlands (BBMRI-NL) [38] and Europe [39] will allow for such studies.

LifeLines DEEP is also, by itself, a unique data source. For 1500 individuals, we will be able to construct genetic risk profiles for predisposition to many common diseases based on genome-wide association data. Next, we will be able to link these risk profiles to phenotype information, as well as clinical, immunological and other parameters. Using this information, we may already be able to compare high-risk individuals with and without disease complaints to generate hypotheses on resilient individuals. At the same time, LifeLines DEEP also allows for integration across different data levels to study, for example, the association between molecular and phenotypic data to increase our understanding of pathogenic mechanisms[40].

One area of particular interest is the domain related to gastrointestinal health. We therefore studied the prevalence of IBS in LifeLines DEEP. We identified IBS in 21% of the participants. These data should be interpreted with caution as our diagnosis is based solely on a participant-administered Rome III criteria questionnaire, and results therefore may be slightly inflated. Nevertheless, our result is consistent with previous suggestions that almost one quarter of the population encounters irritable bowel symptoms over the course of their lifetimes.[41] The prevalence in our cohort confirms that IBS is a common disease and thus research aimed on improved diagnosis and treatment will serve the society. Gastrointestinal complaints are multifactorial making large cohorts necessary to study them in more detail. For IBS in particular there is an urgent need to develop biomarkers that are predictive of the disease. We have selected six gastrointestinal-health-related biomarkers that are currently being analyzed. In a next phase, we will combine our results with other cohorts that study gut health and IBS including the Maastricht IBS cohort (currently including 400 cases and 200 healthy controls, recruitment is ongoing) from the Netherlands, and the TwinGene (n=11,000) and LifeGene (n=30,000, recruitment is ongoing) cohorts from Sweden.

Despite the high prevalence of IBS, the quality of life in our study population in general was higher compared to a random selection of the Dutch population as reported by Aaronson et al.[42] This might be due to age and gender differences, since the National sample included 56% men with a mean(sd) age of 47.6(18.0) years[42], compared to 41% men with a mean(sd) age of 44.6(13.8) years in the LifeLines DEEP cohort. Secular changes may also play a role, since the national survey was conducted more than 15 years ago.

In conclusion, we have established a cohort whose multiple data layers allow for integrative analysis of populations, for translation of this information into biomarkers for disease and for providing new insights into disease mechanisms and prevention.

## ACKNOWLEDGEMENTS

We would like to thank the LifeLines participants and the personnel from the LifeLines study site Groningen for the collaboration. In addition, we would like to thank the LifeLines DEEP research assistants, Wilma Westerhuis-van der Tuuk, Marc Jan Bonder, Astrid Maatman, Mathieu Platteel, Kim de Lange and Debbie van Dussen for their practical and analytical work. We kindly acknowledge Jackie Senior and Kate Mc Intyre for editing our manuscript. Furthermore, we would like to thank The Target project (http://www.rug.nl/target) for providing the compute infrastructure and the BigGrid/eBioGrid project (http://www.ebiogrid.nl) for sponsoring the imputation pipeline implementation.

This project was funded by Top Institute Food and Nutrition Wageningen GH001 to CW, the Biobanking and Biomolecular Research Infrastructure Netherlands (BBMRI-NL) RP3 to LF and an ERC advanced grant ERC-671274 to CW. Sasha Zhernakova holds a Rosalind Franklin fellowship (University of Groningen).

## ETHICAL STATEMENTS

The LifeLines DEEP study has been approved by the ethical committee of the University Medical Centre Groningen, document no. METC UMCG LLDEEP: M12.113965. All participants signed their informed consent prior to study enrolment.

## SUPPLEMENTARY

**Table 3.** Mean and sd of crude and adjusted quality of life scores, two component scores and eight group scores in the LifeLines DEEP population (n=1539) compared to a national sample of the Dutch population [34,42]

| Quality of life |  | Crude | Adjusted<sup>1</sup> | National (n=1742) |
| --- | --- | --- | --- | --- |
| Physical Component Score | mean(sd) | 53.6(7.0) | 53.7(7.0) | 49.7(9.3)[34] |
| Mental Component Score | mean(sd) | 50.4(7.9) | 50.5(7.8) | 52.1(9.6)[34] |
| Physical Functioning | mean(sd) | 92.1(12.3) | 92.2(12.1) | 83.0(22.8)[42] |
| Role-Physical | mean(sd) | 88.5(27.6) | 88.6(27.5) | 76.4(36.3)[42] |
| Bodily Pain | mean(sd) | 85.1(18.2) | 85.2(18.1) | 74.9(23.4)[42] |
| General Health | mean(sd) | 72.8(16.8) | 72.9(16.3) | 70.7(20.7)[42] |
| Vitality | mean(sd) | 67.2(15.5) | 67.4(15.1) | 68.6(19.3)[42] |
| Social Functioning | mean(sd) | 88.3(16.6) | 88.4(16.4) | 84.0(22.4)[42] |
| Role-Emotional | mean(sd) | 90.9(24.7) | 91.3(24.1) | 82.3(32.9)[42] |
| Mental Health | mean(sd) | 79.3(12.6) | 79.4(12.4) | 76.8(17.4)[42] |

<sup>1</sup> adjusted for gender and age

